# Evaluating the Effect of Semantic Congruency and Valence on Multisensory Integration

**DOI:** 10.1101/2021.07.28.454034

**Authors:** Elyse Letts, Aysha Basharat, Michael Barnett-Cowan

## Abstract

Previous studies demonstrate that semantics, the higher level meaning of multi-modal stimuli, can impact multisensory integration. Valence, an affective response to images, has not yet been tested in non-priming response time (RT) or temporal order judgement (TOJ) tasks. This study aims to investigate both semantic congruency and valence of non-speech audiovisual stimuli on multisensory integration via RT and TOJ tasks (assessing processing speed (RT), point of subjective simultaneity (PSS), and time-window when multisensory stimuli are likely to be perceived as simultaneous (Temporal Binding Window; TBW)). Forty participants (mean age: 26.25; females=17) were recruited from Prolific Academic resulting in 37 complete datasets. Both congruence and valence have a significant main effect on RT (congruent and high valence decrease RT) as well as an interaction effect (congruent/high valence condition being significantly faster than all others). For TOJ, images high in valence require visual stimuli to be presented significantly earlier than auditory stimuli in order for the audio and visual stimuli to be perceived as simultaneous. Further, a significant interaction effect of congruence and valence on the PSS revealed that the congruent/high valence condition was significantly earlier than all other conditions. A subsequent analysis shows there is a positive correlation between the TBW width (*b*-values) and RT (as the TBW widens, the RT increases) for the categories that differed most from 0 in their PSS (Congruent/High and Incongruent/Low). This study provides new evidence that supports previous research on semantic congruency and presents a novel incorporation of valence into behavioural responses.

## Introduction

In our everyday environment, we are constantly exposed to a barrage of multi-modal stimuli (i.e., images, sounds, smells, etc.). Cross-modal, multisensory integration is a complex process which allows for production of a coherent percept (Spilcke-Liss et al., 2019). Frequently, surrounding stimuli can be conflicting; therefore, understanding how the central nervous system (CNS) is able to resolve conflicts of sensory inputs when making judgements is a large area of study. The binding of separate stimuli (multisensory integration) is known to be impacted by factors such as spatial and temporal proximity and has been more recently shown to be impacted by the semantic (higher level meaning of stimuli) congruency of the stimuli (Barutchu et al., 2018; Delong & Noppeney, 2021; Doehrmann & Naumer, 2008; Spilcke-Liss et al., 2019). Being able to understand how different factors, such as semantics and valence (affective response to a stimulus), can modulate this resolution of conflicting stimuli is key to understanding multisensory integration. This semantic congruency can take many forms and can be evaluated through response time, object identification, and judgements of temporal order and synchronicity. By pairing congruent and incongruent stimuli of different levels of valence we can begin to better understand the processes the brain undertakes to integrate the stimuli in our surroundings.

### Multisensory integration and sensory conflict

Multisensory integration is the ability of the brain to synthesize information from two or more different senses into a coherent output/percept (Stein et al., 2002; Stein & Stanford, 2008). Inputs to the brain can be either unimodal (inputs originating from only one sense) or cross-modal (inputs from two or more different senses) (Stein et al., 2002; Stein & Stanford, 2008). Multisensory integration takes place even when the inputs from the modalities are in conflict and this resulting percept is a frequent area of research (Stein & Stanford, 2008). There are a number of well-known effects of multisensory integration including the McGurk Effect and the ventriloquism effect (see Delong & Noppeney, 2021; Laurienti et al., 2004; Stein & Stanford, 2008). These effects alter our perception of an event from the “true” stimuli. The McGurk effect alters the understood meaning of speech stimuli, while the ventriloquism effect alters the perceived location of an auditory stimulus (Howard & Templeton, 1966 as cited in Laurienti et al., 2004; McGurk & MacDonald, 1976; Stein & Stanford, 2008). These examples of cross-modal effects and illusions help to enhance the understanding of multisensory integration in the brain, but leave more questions to be explored about other factors that impact multisensory integration, as done in this study.

### Behavioural Indicators of Multisensory Integration

A frequently utilized behavioural measure of multisensory integration is response time (RT) (Colonius & Diederich, 2017). RT is often measured by the time it takes a participant to respond (typically via a button press) to the appearance of a stimulus or multiple stimuli. Important to note here is the use of response time versus reaction time: while the term reaction time is often used in related literature, true reaction time is defined as, “a measure of the time from the […] signal to the *beginning* of the response to it” (Schmidt, 1988, p. 64). Response time, alternatively, is defined as, “the sum [of] RT [reaction time] and MT [movement time]” where movement time is the “interval from the initiation of the response (the end of RT [reaction time]) to the completion of the movement” (Schmidt, 1988, p. 65). Thus, a true response time is the time from the initial signal to the completion of the movement (Schmidt, 1988). As such, in this study, the term response time will be used as it the most accurate for this task. Examples of different response time tasks include stimuli categorization into colour (Laurienti et al., 2004), animacy (Steinweg & Mast, 2017), and plausibility (Spilcke-Liss et al., 2019). When measuring multisensory integration, the time to respond to cross-modal stimuli is usually shorter than the RT of any unimodal stimulus (Colonius & Diederich, 2017; Laurienti et al., 2004; Steinweg & Mast, 2017).

Other frequently used measures of multisensory integration include temporal order judgement (TOJ) and simultaneity judgement (SJ) tasks. In both tasks, two cross-modal stimuli are presented either synchronously or asynchronously, with varied timings of onset (Stimulus Onset Asynchrony; SOA) (Basharat et al., 2018; Love et al., 2013). In the TOJ task, participants are asked to determine which stimuli was presented first, while in the SJ task they are asked to determine whether the stimuli were synchronous or not (Basharat et al., 2018; Love et al., 2013). Both tasks can measure participant accuracy (Point of Subjective Simultaneity; PSS) and variance (Just Noticeable Difference; JND), however, the tasks themselves often differ in results, thus showing that they may evaluate different perceptual processes (Basharat et al., 2018; Bedard & Barnett-Cowan, 2016; Love et al., 2013).

### Semantics

A more recent area of research is investigating semantics on cross-modal, multisensory integration. Semantics can indicate a wide variety of manipulations but are generally categorized as manipulations of more complex and higher level variables, frequently the meaning and context of the stimuli (Doehrmann & Naumer, 2008; Laurienti et al., 2004). Studies evaluating semantics often make use of and manipulate the network of information collected in semantic memory to test congruent (i.e., an image of a cat and a matching meow sound) and incongruent (an image of a cat and a non-matching bark sound) conditions. Most commonly, the cross-modal stimuli are auditory and visual stimuli, but some investigations into other modes do exist (for example, Gottfried & Dolan (2003) used olfaction). These investigations have indicated that semantic congruency/incongruency impacts a number of the behavioural effects (such as response time and temporal order judgements) of multisensory integration (Cox & Hong, 2015; Doehrmann & Naumer, 2008; Laurienti et al., 2004).

### Semantic Congruence and Multisensory Integration

Laurienti and colleagues (2004) were among the first to demonstrate higher level semantic effects on cross-modal multisensory integration. Using audio-visual (AV) stimuli of coloured disks with written or spoken colour names, they demonstrated that cross-modal (auditory and visual) semantically congruent stimuli produced faster response times compared to incongruent or unimodal stimuli (Laurienti et al., 2004). The finding of faster congruent speed responses was only found for the cross-modal stimuli and not the dual-visual condition (coloured circle and written colour name) although incongruent stimuli increased response time for both the dual-visual and cross-modal conditions. This demonstrates that the semantic content of stimuli does in fact impact multisensory integration and the resulting behaviours, as will be further investigated in this study.

A study by Vatakis and Spence (2007) investigated the “unity assumption” and its relation to cross-modal, multisensory integration. The “unity assumption” can be defined as, “the assumption that a perceiver makes about whether he or she is observing a single multisensory event rather than multiple separate unimodal events” (Vatakis & Spence, 2007, p. 744). While the “unity assumption” and semantic congruency are similar (both are top-down modulatory factors in multisensory integration), they are not exactly the same concepts and may not have identical effects on multisensory integration (see fig. 1) (Chen & Spence, 2017). More thoroughly discussed in Chen & Spence (2017), it can be difficult to separate differences between semantic congruency and the unity assumption. Through the use of auditory words and syllables presented with videos of faces uttering the syllables and words, Vatakis and Spence (2007) demonstrated that in the “matched” or congruent cases (where the audio presented matched the video clip lip movements), participants had more difficulty determining temporal order compared to the “mismatched” or incongruent cases. This shows evidence that in the congruent cases, participants made the assumption that the two stimuli were from one event which enhanced the multisensory binding (Vatakis & Spence, 2007). This shows that the “unity assumption”, as well as higher level semantic meaning, can modulate cross-modal, multisensory integration.

**Figure 1.**
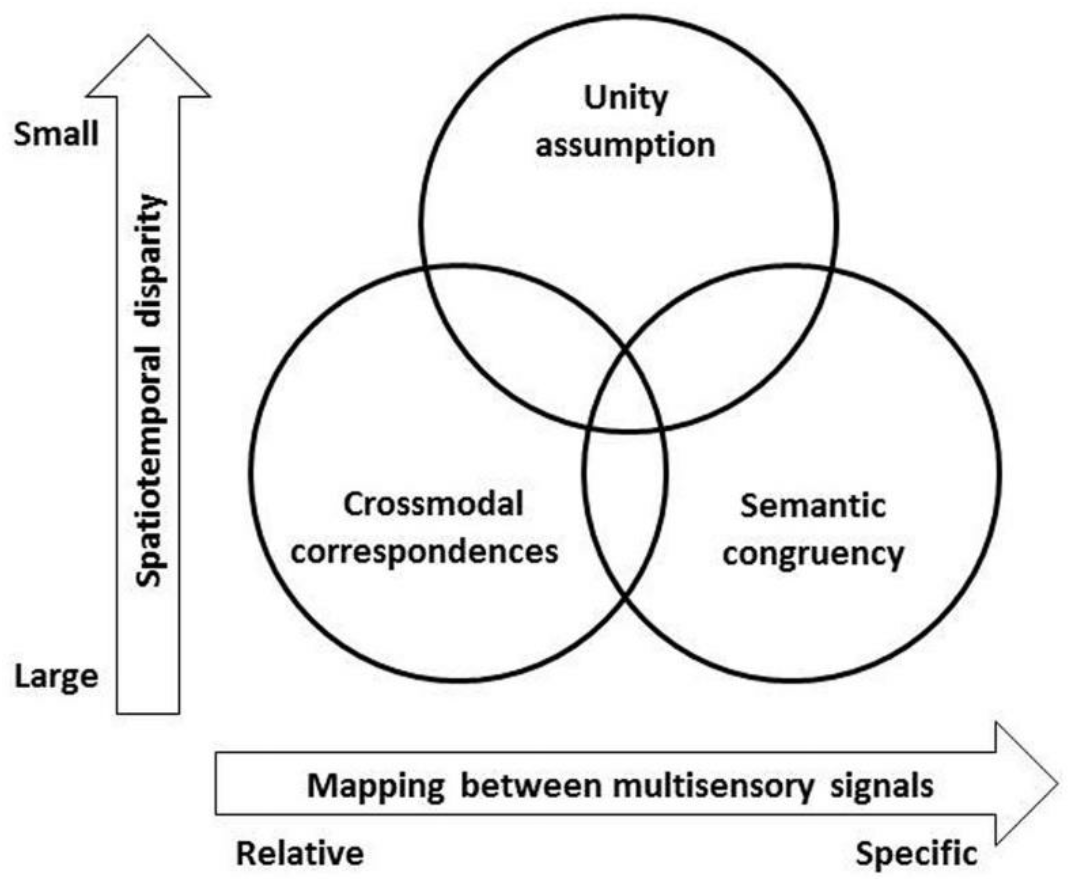
Schematic to represent relationships between modulatory factors in MSI. (Chen & Spence, 2017)

Following their 2007 study, Vatakis and Spence (2008) attempted to replicate their results using non-speech stimuli (videos of actions with the corresponding sounds). They were unable to produce the same “unity assumption” with non-speech stimuli, perhaps due to the nature of the stimuli. Important to note here is that semantic congruency does differ from the “unity assumption” (see fig. 1) and non-speech stimuli have been repeatedly shown to impact the behavioural effects of cross-modal multisensory integration in other studies (Chen & Spence, 2017; Cox & Hong, 2015; Spilcke-Liss et al., 2019).

A further study investigating semantic effects of non-speech stimuli (Chen & Spence, 2010) used a variety of line drawings of animals and objects that were presented very briefly and with either a congruent or incongruent sound (at different SOAs) to investigate semantic effects on image identification. They found that semantically congruent sounds aided in image identification (as long as the audio was presented within at least 300 ms of the image) compared to white noise or an absence of noise, while semantically incongruent audio worsened image identification (Chen & Spence, 2010). This suggests that semantic meanings associated with the auditory and visual stimuli are processed in an overlapping semantic system, and that congruence between them can aid multisensory integration (Chen & Spence, 2010).

Steinweg and Mast (2017) investigated semantic congruency of both colours with spoken words and animals/objects with their sounds. In experiment 1, participants were asked to press a button if two of the four colours were presented either aurally or visually and a different button for the other two colours (Steinweg & Mast, 2017). In experiment 2, the stimuli were images and sounds of animals and objects, and in each trial participants were asked to decide as quickly as possible if the stimuli presented were animate or inanimate (Steinweg & Mast, 2017). Response time and error rates were collected in both experiments, and overall results showed fewest errors in the incongruent condition and the fastest response time with the congruent condition (Steinweg & Mast, 2017). The bimodal stimuli (congruent and incongruent) had faster response times than the unimodal stimuli, and the semantically congruent stimuli had faster response times than the semantically incongruent stimuli. They were able to show that the semantically incongruent condition increased response caution (via reduced error rates and using the drift-diffusion models which investigate the speed-accuracy trade-off) which led to the increased response time compared to the congruent condition (Steinweg & Mast, 2017). Thus, semantic congruency has an impact on the behaviour effects of multisensory integration. This idea is further investigated in the present study.

When investigating multisensory integration, it is also important to consider attentional effects, as done by Spilcke-Liss and colleagues (2019). They asked participants to attend to two of three possible stimuli (visual foreground, visual background, auditory stimuli) and determine if those two were congruent (“plausible”) or not and recorded error rate and response times. They opted to ask “plausible” rather than congruent as participants understood “plausible” better than “congruent” (Spilcke-Liss et al., 2019). They found that the error rates were significantly higher when the unattended stimulus was incongruent compared to the two attended and congruent stimuli (Spilcke-Liss et al., 2019). This shows that semantic incongruency, even in unattended stimuli, can disrupt perceptual processing of stimuli. Additionally, they found that response time results did not follow the pattern of errors (no significant difference in times between correct and incorrect trials) and appeared to be unrelated to the attentional manipulation (Spilcke-Liss et al., 2019). They discussed that semantic congruency impacts multisensory integration via non-voluntary attentional increases due to congruent (speeded processing) and incongruent (disrupted processing) stimuli (Spilcke-Liss et al., 2019). Thus, even if not endogenously attended, semantic congruencies have an impact on multisensory integration in our perceptual system. Interestingly, Cox and Hong (2015) further demonstrated that not only do the stimuli not have to be attended for semantic congruency to modulate multisensory integration, the individual does not even have to be aware of the visual stimuli for semantics facilitated multisensory integration to take place.

More recently, Delong and Noppeney (2021) investigated the impact and interaction of both sematic congruency and spatial correspondences. They conducted two experiments using realistic images and sounds as well as masking in experiment 2. In each trial participants were presented with one or two images that varied in location on the screen and either a semantically congruent or incongruent sound from either the right or left side (or bilaterally) (Delong & Noppeney, 2021). In both experiments, participants were asked to identify where the sound came from (left or right) and in experiment 2 they were additionally asked to identify which image(s) they saw. In experiment 1, they found that semantically congruent stimuli significantly improved audiovisual binding and increased the impact of spatial congruence (Delong & Noppeney, 2021). Additionally, there was an interaction effect between the semantic and spatial congruences where the effect of the semantic congruence on the audiovisual binding was increased when the sound was presented bilaterally (Delong & Noppeney, 2021). In experiment 2, they found a significant positive effect of semantically congruent stimuli on picture identification (in line with previous studies such as Chen and Spence (2010)). Experiment 2 also found a significant interaction effect of semantic and spatial congruences (as in experiment 1) but only for the visible conditions (conditions where the participant identified that they saw and recognized the image) as opposed to the invisible conditions (where the participants reported that they could not identify the image that was presented) (Delong & Noppeney, 2021). Ultimately, this study further reinforces the impact that semantic congruency has on our multisensory integration and demonstrates that it is related to other factors such as spatial congruency. This study builds on this semantics interaction by investing its relationship with valence.

A number of studies have investigated the effect of semantic congruence on multisensory integration as is described here (a summary can be found in Table 1 below). These studies provide context and the motivation for the further investigation of semantic congruency and multisensory integration examined in this study.

**Table 1.** Summary of studies investigating semantic congruency and multisensory integration

| Study | Concept investigated | Test used | Effect of semantic congruency on multisensory integration? |
| --- | --- | --- | --- |
| Laurienti et al (2004) | Semantic congruency | RT and error rate via feature discrimination | Yes |
| Vatakis & Spence (2007) | Unity assumption | TOJ | Yes |
| Vatakis & Spence (2008) | Unity assumption | TOJ | No |
| Chen and Spence (2010) | Semantic congruency | Image identification | Yes |
| Cox & Hong (2015) | Semantic congruency | RT and visual event identification | Yes |
| Steinweg & Mast (2017) | Semantic congruency | RT and error rate via stimulus categorization | Yes |
| Spilcke-Liss et al. (2019) | Semantic congruency and endogenous attention | RT and error rate via stimulus categorization | Yes |
| DeLong & Noppeney (2021) | Semantic and spatial congruency | Image identification and sound localization | Yes |

### Valence

When presented with an image, emotional processes are activated which have been shown to impact perceptual processing and modulate attention (Gerdes et al., 2014). Additionally, there is evidence of cross-modal influences of emotion. Thus, when using images as stimuli in research, it is important to evaluate and control for the affective responses from the images. Two common affective evaluations of images are valence (positive or negative affective response) and arousal (level of excitement experienced by an observer) (Gerdes et al., 2014; Kurdi et al., 2017). In a study of affective priming using words, valence was shown to have a differential and stronger effect on target processing than arousal as well as being more stable than arousal in the semantic system (Yao et al., 2019). Additionally, a further affective priming study (using naturalistic auditory stimuli) showed that arousal did not influence the priming effects while valence did (Topolinski & Deutsch, 2013). As discussed in Yao et al. (2019), positive valence has been shown in some cases to have a faster response time (Yao & Wang, 2013). As it relates to semantics, studies discuss affective processes as contributing to and/or being a category of semantics (Avero & Calvo, 2006; Topolinski & Deutsch, 2013). Despite this, in studies of semantic congruency, valence has not been investigated and it is unclear whether valence effects seen in priming studies extend to other tasks such as RT and TOJ. As such, the present study will incorporate the affective valence ratings of images as a semantic category (see fig. 2).

**Figure 2.**
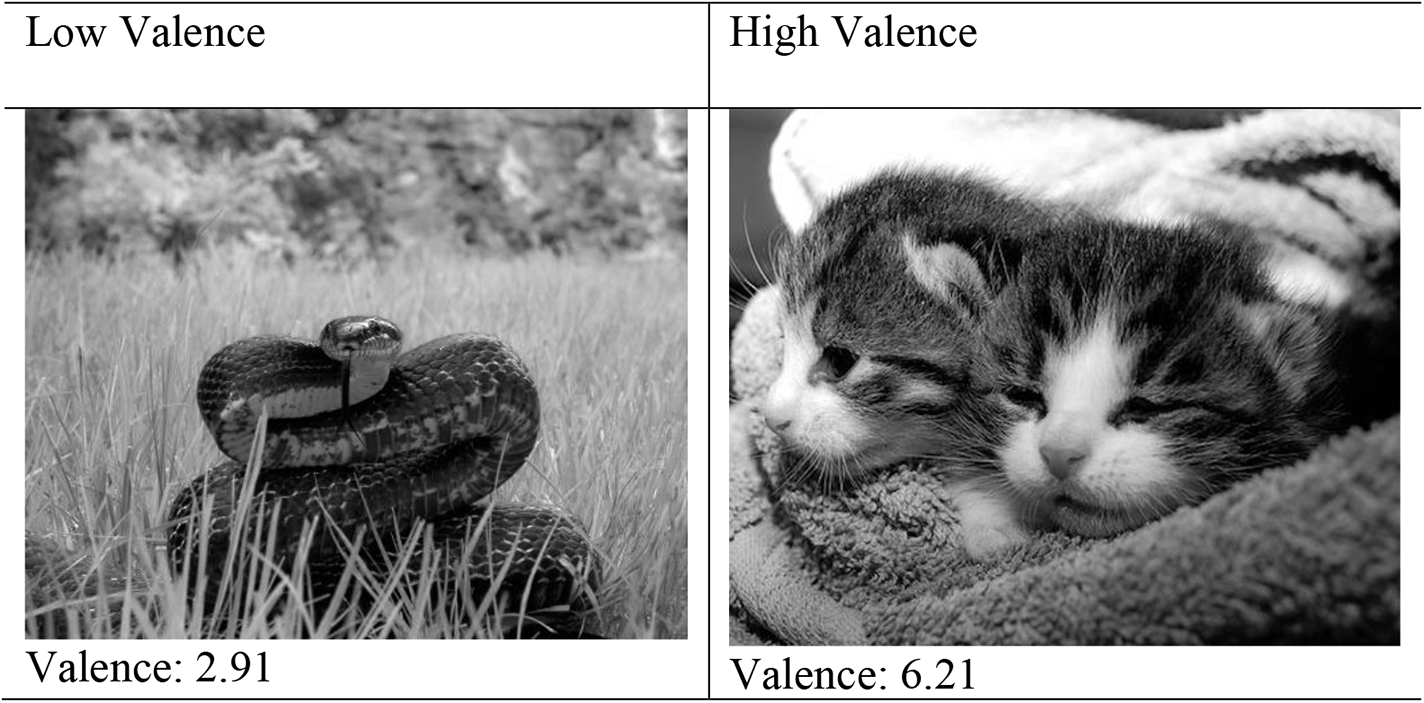
Example images of low and high valence. Images from OASIS, where valence is rated from 1 (very negative) to 7 (very positive). Original images are in colour but black and white versions were used for this study (Kurdi et al., 2017).

To incorporate the valence of images into research, the valence of the images has to be first determined. Data sets of images with pre-determined valences do exist, such as the International Affective Picture System (IAPS) and Open Affective Standardized Image Set (OASIS) (Bradley & Lang, 2017; Kurdi et al., 2017). The OASIS set is used in this study due to its open-source nature. Researchers, after explaining the terms valence and arousal, asked participants to rank the 900 images for these two attributes (Kurdi et al., 2017). With over 800 responses, they determined the average valence and arousal for each individual image, thus creating the OASIS set (Kurdi et al., 2017).

### Animacy

Another category that semantics can be divided into is animacy categories (animate and inanimate) (Praß et al., 2013). As discussed in Praß and colleagues (2013), different processing mechanisms for animate vs inanimate stimuli have been suggested by a number of studies, yet there is no clear indication of a behavioural advantage for either category. Some studies have shown the animate category to have a faster response time compared to the inanimate category (see Crouzet et al., 2012) while others have shown the opposite (such as Praß et al., 2013). As it is unclear whether animacy has an effect, in this study, we decided to use only animate stimuli to avoid any confounding factors.

### Research Question and Hypothesis

This study aims to answer the following research questions:

1. Does semantic congruency of non-speech audiovisual stimuli impact multisensory integration as seen through RT and TOJ?
2. Does the valence of non-speech audiovisual stimuli impact multisensory integration as seen through RT and TOJ?
3. Do congruency and valence interact to impact multisensory integration as seen through RT and TOJ?

Based on the literature, the main objective of the present study is to investigate the impact of semantic congruency and valence on the behavioural effects (RT and TOJ) of multisensory integration. We hypothesize that (1) the congruent conditions will have faster response times and worse ability to discriminate temporal order, (2) positive valence will lead to a faster response time and may impact temporal order judgements and (3) that valence and congruence may interact (positive valence and semantically congruent stimuli will have the fastest response time).

## Methods

### Participants

Participants were recruited and tested through Prolific Academic. Inclusion criteria included participants being 18+ years of age with normal or corrected-to-normal vision and hearing. Participants were remunerated at a rate of $10 CAD/hour for their participation in this study. This study received approval from the University of Waterloo’s Human Research Ethics Board. Written informed consent was obtained from all participants before participation. A sample size power calculation was conducted using G*Power and provided a required sample size of 24. Given the fact that online studies are known to have lower quality data, the expected sample size was increased to 40 to ensure valid data of at least 24 were obtained.

### Experimental set-up and stimuli

The experiment was coded using PsychoPy version 2020.2.8 in builder mode (https://www.psychopy.org/), hosted and run on Pavlovia (https://pavlovia.org/), and recruitment was conducted through Prolific Academic (https://www.prolific.co/). Data from this study is available in the Open Science Framework (https://osf.io/atkfq/ ; DOI 10.17605/OSF.IO/ATKFQ). Participants used their own personal computing device at an arm’s length distance (as per Bechlivanidis & Lagnado, 2016). In all tasks, a fixation cross (of combined bulls-eye and cross hair design) was presented prior to and following the visual stimuli presentation (as recommended by Thaler et al., 2013). The visual stimuli (500 x 400 pixels) were grayscale photos adapted from the OASIS database (Kurdi et al., 2017). Fixation crosses were presented for 1000 ms and after a varying delay of 0-500 ms (to reduce temporal predictability), the visual stimulus was presented for 500 ms, followed again by the fixation cross. The fixation cross and visual stimuli were presented in the centre of the screen. Auditory stimuli (that were typical sounds of each of the visual stimuli) were retrieved from web sources (i.e., from www.findsounds.com) and were presented for 500 ms. Participants were instructed to have device volume at maximum comfortable level. Auditory stimuli were played through the computing device itself (thus the dB is unknown for each participant, but participants set the volume such that it was above detection threshold). Responses were made by keyboard button press. Participants had to make a response to complete a trial and progress to the next. A 3000 ms response window was provided and, should they have failed to respond within this time, that trial was failed and was presented again later in the task.

### Design and procedure

The experiment is composed of two tasks: temporal order judgement and response time. Each participant proceeded through the trials in a randomized order. The RT task and TOJ task consist of 416 trials each, and with approximately 3-4s per trial, the entire study took approximately 45 minutes for each participant to complete.

#### RT Task

The RT task has 4 image/sound combinations (see Table 2) and has 104 repetitions of each image/sound combination for a total of 416 trials. Prior to the 416 trials, participants have 6 practice trials in order to become familiar with the task (Bedard & Barnett-Cowan, 2016). Participants are presented with the image and auditory stimuli simultaneously and are asked to make a decision as to whether the two are “plausible” or not, an accessible way of asking congruency (Spilcke-Liss et al., 2019). They communicate this with a button press where “1” represents “plausible” and “2” represents “not plausible”. Participants are instructed to respond as quickly and correctly as possible.

**Table 2.**
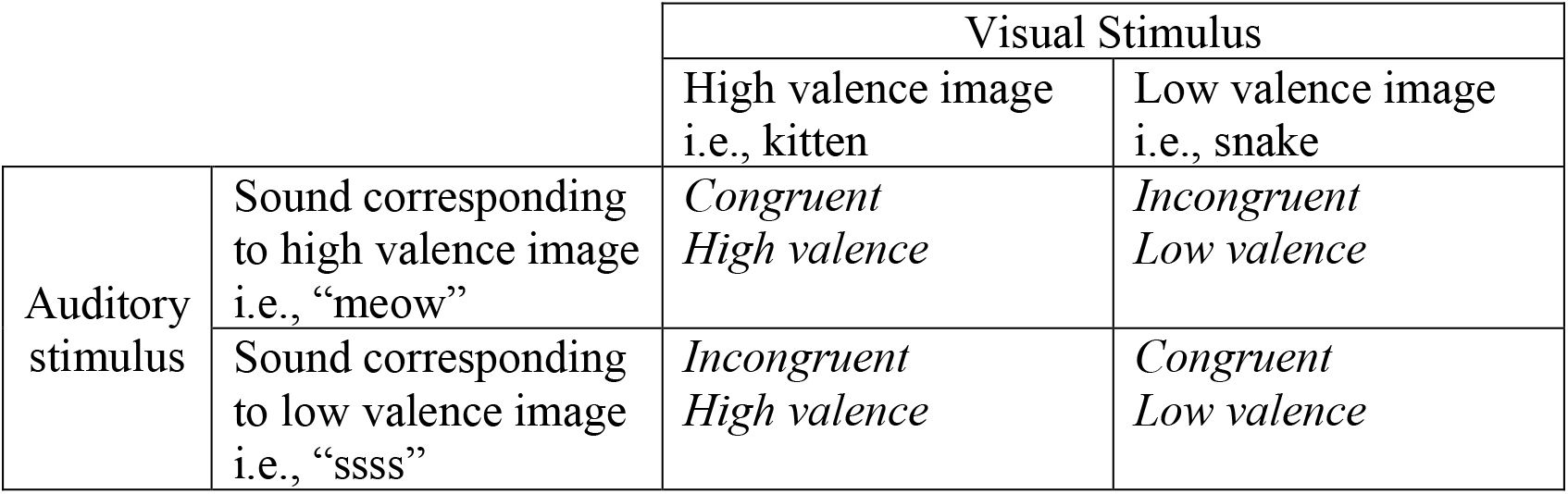
The four categories of image/sound conditions by congruency and valence

|  |  | Visual Stimulus |  |
| --- | --- | --- | --- |
|  |  | High valence image<br>i.e., kitten | Low valence image<br>i.e., snake |
| Auditory<br>stimulus | Sound corresponding<br>to high valence image<br>i.e., “meow” | *Congruent*<br>*High valence* | *Incongruent*<br>*Low valence* |
|  | Sound corresponding<br>to low valence image<br>i.e., “ssss” | *Incongruent*<br>*High valence* | *Congruent*<br>*Low valence* |

#### TOJ Task

The TOJ task has 4 image/sound combinations (see Table 2), 13 stimulus onset asynchronies (SOAs), and 8 trials of each, resulting a total of 416 trials. Prior to the testing trials, participants have 6 practice trials to become familiar with the task (Bedard & Barnett-Cowan, 2016). The SOAs are ±300, ±200, ±150, ±100, ±50, ±25, and 0 ms, where negative values indicate that the auditory stimulus was presented first (Basharat et al., 2019; Bedard & Barnett-Cowan, 2016). Participants are presented with both visual and auditory stimuli in all trials, but they are presented at each of the SOAs in random order. Participants are asked to decide whether the image (“3” button press) or sound (“4” button press) was presented first. Participants are asked to respond the most correctly.

### Statistical Analysis Methods

This study is a within-subjects design and the tasks (RT and TOJ) were counter-balanced. The independent variables are congruency (levels are congruent and incongruent) and valence (levels are high and low), and the dependent variables are RT and TOJ outputs (RT outputs are mean and median response times as well as accuracy and TOJ outputs are the PSS and JND). The data collected from each task (RT and TOJ) was analyzed separately using JASP 14.1.0. The RT data analyzes main effects of congruency and valence as well as interaction effects with a 2×2 repeated measures ANOVA. The data from the TOJ task evaluates main effects of semantic congruency and valence as well as interaction effects in 2 separate 2×2 repeated measure ANOVAs, one for the PSS and one for the JND.

The accuracy (PSS values) and precision (JND) of participant judgements in the TOJ task were estimated using SigmaPlot version 12.5. Psychometric functions were fitted to responses as a function of SOA. Analysis was done individually for each participant using a sigmoidal logistic regression:

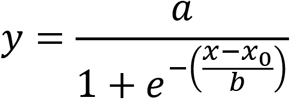

Where *a* is the amplitude (fixed to 1 in the function), *x_0_* is the PSS, and *b* is the slope (and acts as a proxy for the JND).

## Results

### Participants

A total of 40 Prolific participants completed the study. Of these 40, the mean age is 26.25 (range: 18-57), 17 identify as female, and 23 identify as male. Three participants encountered technical errors and thus a complete data set is available for 37 participants. Of the 37 complete, 1 had low quality RT data (each category less than 70% accuracy) and 10 had low quality TOJ data (an R^2^ value of less than 0.2). As such there were 36 complete data sets in the RT analysis and 27 complete data sets in the TOJ analysis, both above the number required in the power analysis.

### Online Results

As with other recent papers, our data obtained from Prolific Academic recruitment was of overall high quality, comparable to in-lab data (Crump et al., 2013; Woods et al., 2015). We ran a small pilot study in-lab and, via observational assessment, found no differences between quality and results of the pilot and Prolific data. As per Crump et al. (2013), collection was completed within hours and our dropout rate was higher than in-lab, but not prohibitive. The response times were in line with other choice response times and TOJ values were as expected, showing that our study design was valid even in an online environment.

### Response Time Analysis

After removal of one participant with low quality data (accuracy from 38-57%), overall accuracy for each group was as follows: Congruent/High: 96%, Congruent/Low: 85%, Incongruent/High: 92%, Incongruent/Low: 95%.

RT data were checked for normality and all were found to be normally distributed (via the Shapiro-Wilk test). The median response time for each category was calculated individually for each participant, then the mean of median response times was calculated for each category (Congruent/High: 0.738 s; Incongruent/High: 0.839 s; Congruent/Low: 0.805 s; Incongruent/Low: 0.832 s). Median values were used due to the known positive skew of RT results, and thus are best quantified for central tendency by their median (Der & Deary, 2017). Results from the 2×2 RM ANOVA indicate a significant main effect of congruence (F = 90.910; *p* < 0.001; ɳ^2^= 0.335), a significant main effect of valence (F = 15.518; *p* < 0.001; ɳ^2^= 0.071), and a significant interaction effect of congruence and valence (F = 19.884; *p* < 0.001; ɳ^2^= 0.110). A post-hoc test with an applied Bonferroni correction (0.05/4=0.0125) showed that the Congruent/High condition is significantly faster than all other conditions (incongruent/high *p* < 0.001; congruent/low *p* < 0.001; incongruent/low *p* < 0.001) and that the Incongruent/High condition is significantly slower than the Congruent/Low (*p* = 0.006) (see fig. 3).

**Figure 3.**
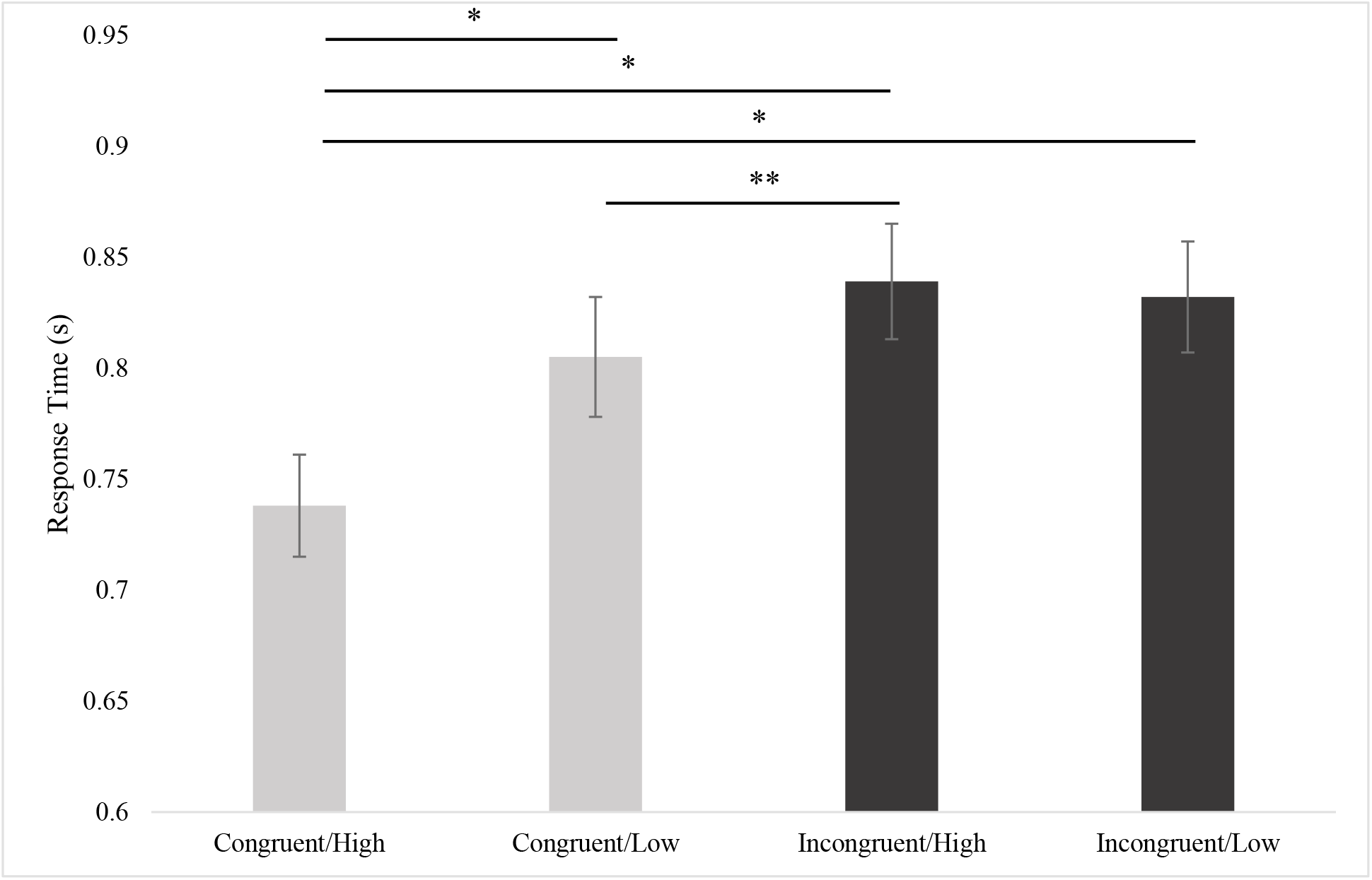
Average median response time 2×2 RM ANOVA results. Average RT values (with SEMs) from each condition. From the Bonferroni post-hoc analysis, a single asterisk (*) indicates *p* < 0.001 and a double asterisk (**) indicates *p* < 0.01.

### Temporal Order Judgement Data Analysis

The individual curves per participant and average curve for each category were calculated and are plotted below in figures 4 (average curves) and 5 (individual and averages by category). The R^2^ was calculated for each fit and data sets with an R^2^ below 0.2 removed from the analysis. There were 10 data sets that were below this cut-off and were thus removed, leaving a total of 27 data sets for analysis.

**Figure 4.**
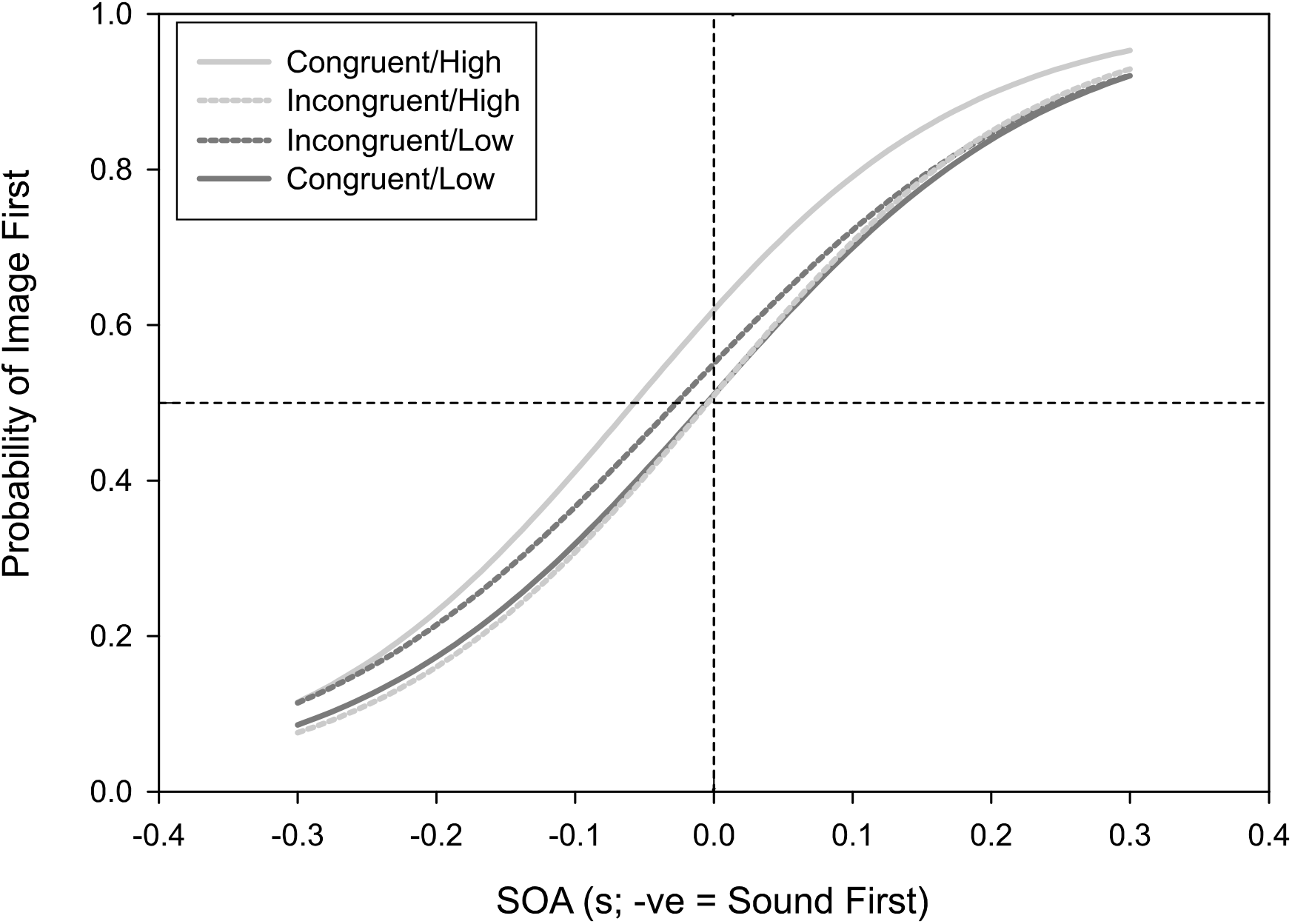
TOJ average curve plots for each category. In order to perceive the two stimuli as simultaneous, the Congruent/High conditions required the **sound** to be presented 0.058 s before the image, the Congruent/Low condition required the **image** to be presented 0.005 s before the sound, the Incongruent/High condition required the **sound** to be presented 0.004 s before the image and the Incongruent/Low condition required the **sound** to be presented 0.027 s before the image.

**Figure 5.**
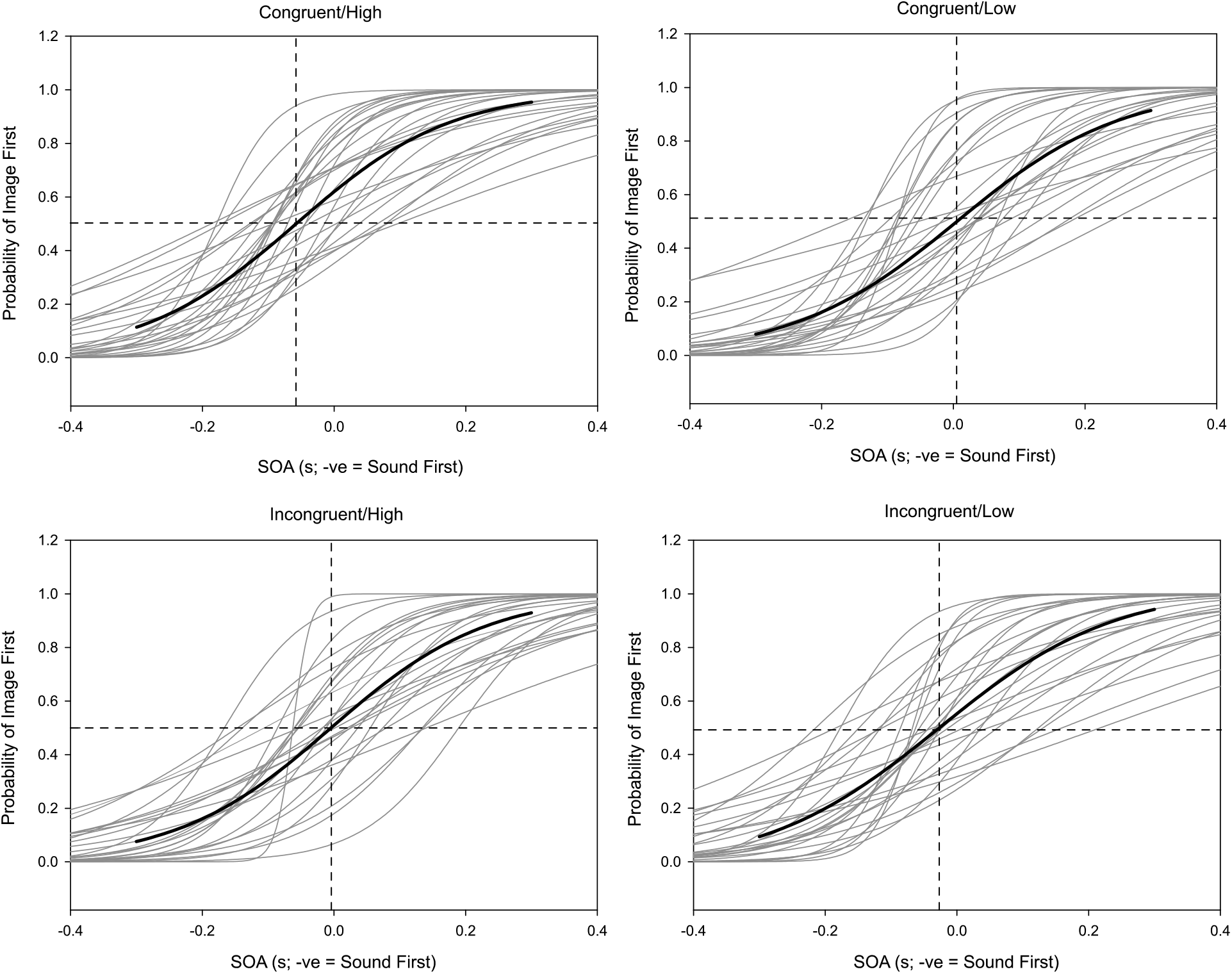
TOJ plots by category with each participant and category average. The sigmoidal functions are fit to average (thick black lines) and individual (thin grey lines) data.

TOJ data were checked for normality and while most were found to be normally distributed (via the Shapiro-Wilk test), the congruent/high, incongruent/low, and congruent/low conditions for the b-values were not normal (Shapiro-Wilk p-values of 0.007, 0.019, and 0.006 respectively). In order to correct for this non-normality, a Box-Cox transformation was applied in R (version 4.0.4) which gave a result of an exponent of 0.1818 (Box & Cox, 1964). All b-values underwent this transformation, after which all data were normally distributed. Separate 2×2 RM ANOVAs were conducted for PSS and JND. For PSS (x_o_), there was no significant main effect of congruence (F = 2.396; *p* = 0.134; ɳ^2^= 0.013), but there was a significant main effect of valence (F = 7.276; *p* = 0.012; ɳ^2^= 0.05) and a significant interaction effect between congruence and valance (F = 15.229; *p* < 0.001; ɳ^2^= 0.226). A post-hoc test with applied Bonferroni correction (0.05/4=0.0125) shows that the Congruent/High condition is lower than all other conditions (incongruent/high (*p* = 0.002), congruent/low (*p* < 0.001), and incongruent/low (*p* = 0.022; see fig. 6).

**Figure 6.**
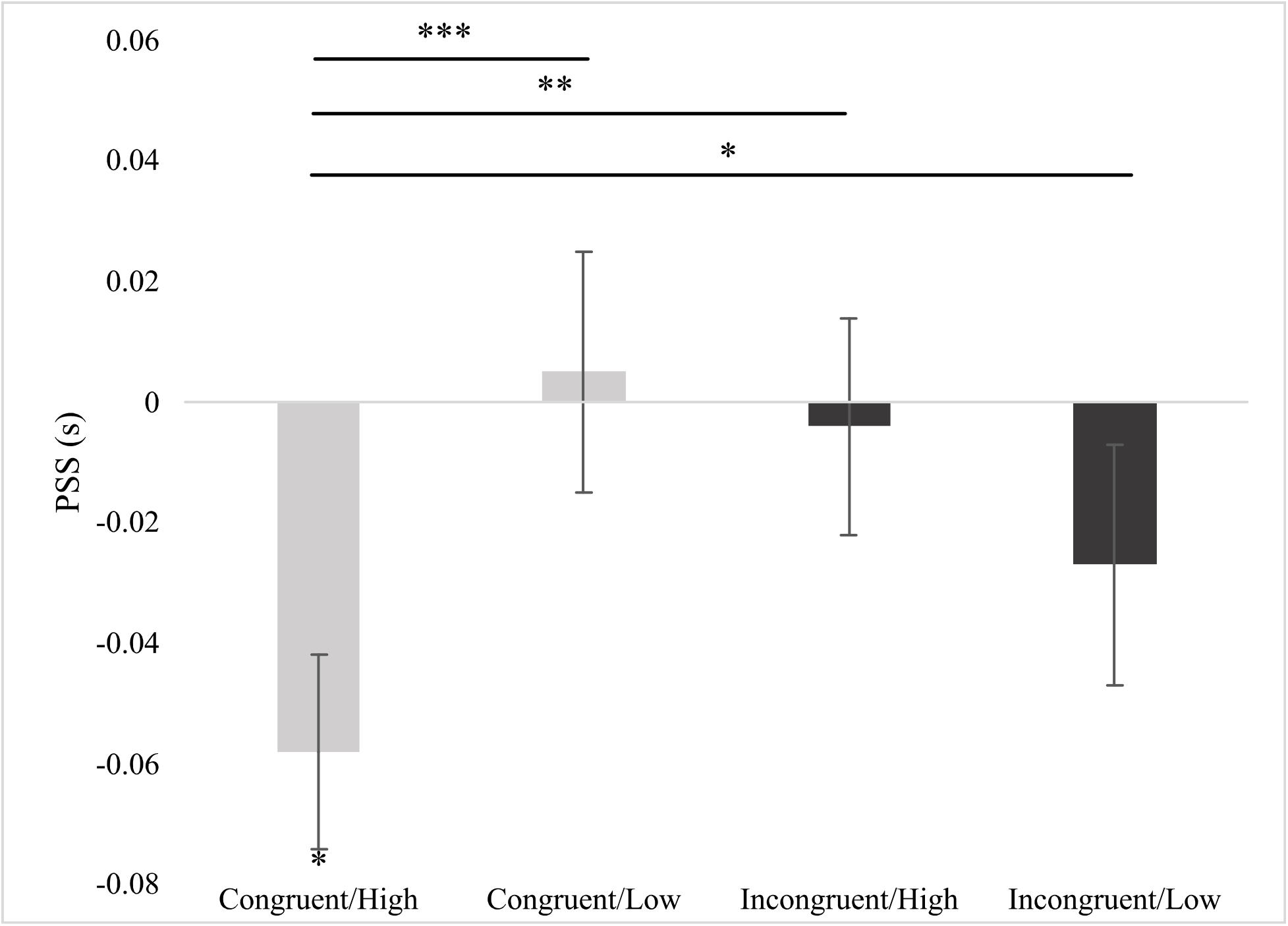
PSS 2×2 RM ANOVA results. Average PSS values (with SEMs) from each condition. From the post-hoc analysis with Bonferroni correction, a single asterisk (*) indicates *p* < 0.05, a double asterisk (**) indicates *p* < 0.01, and a triple asterisk (***) indicates *p* < 0.001. An asterisk near the error bar indicates *p* = 0.001 on the one-sample student T-tests between PSS values and 0. See after figure 7 for full results of t-tests.

For the JND (*b*), prior to the transformation, there was no significant main effect of either congruence (F = 0.373; *p* = 0.547; ɳ^2^= 0.002) or valence (F = 0.946; *p* = 0.340; ɳ^2^= 0.019), and no significant interaction effect (F = 0.215; *p* = 0.647; ɳ^2^= 0.003). After the transformation, there remains no significant main effect of either congruence (F = 0.637; *p* = 0.432; ɳ^2^= 0.005) of valence (F = 0.597; *p* = 0.447; ɳ^2^= 0.009), and no interaction effect (F = 0.183; *p* = 0.673; ɳ^2^= 0.003) (see fig. 7).

**Figure 7.**
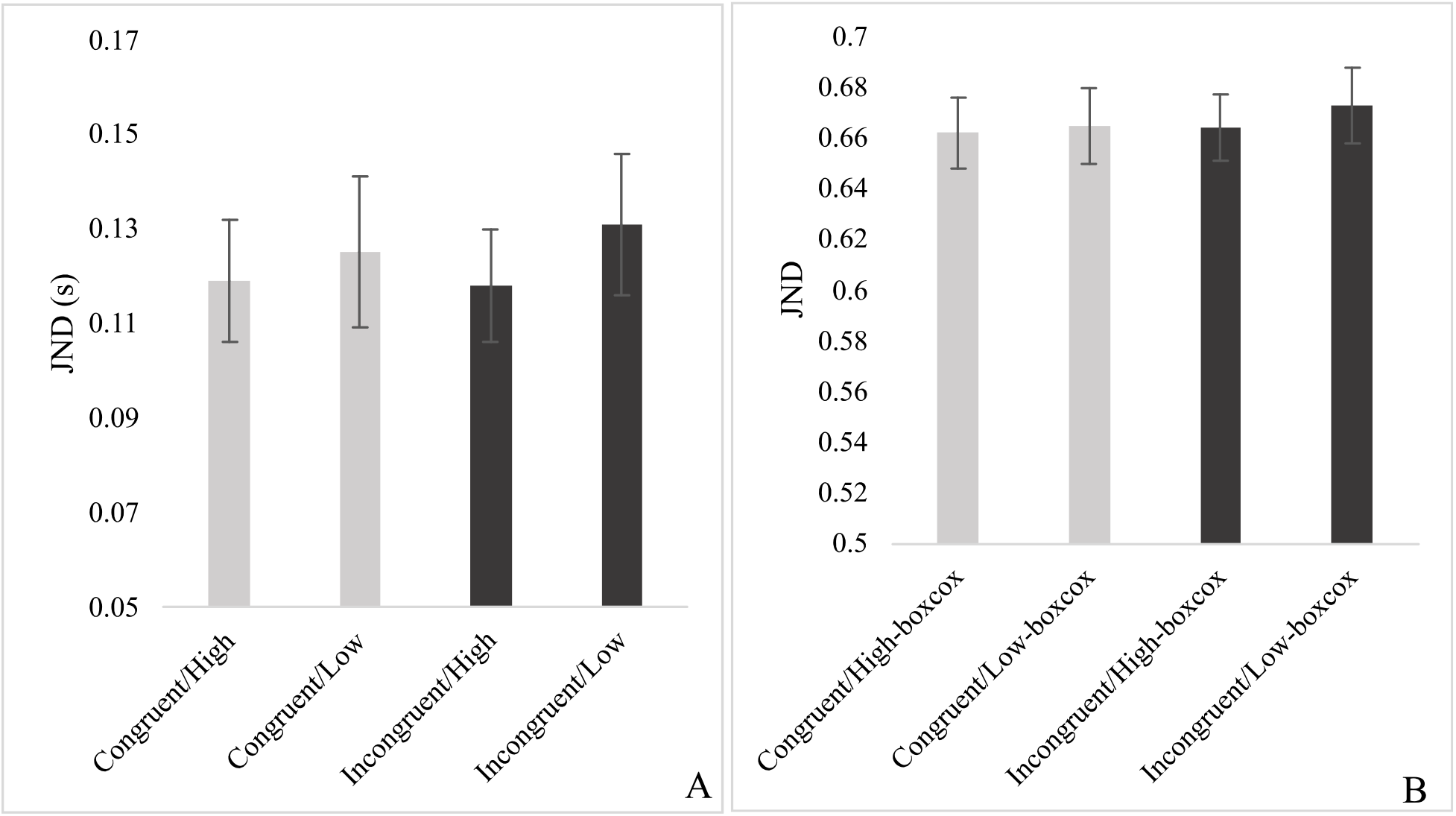
JND 2×2 RM ANOVA results. Panel A reports the true values (with SEMs) and panel B reports the transformed values after normality corrections (with SEMs). In both, there are no significant effects.

Following this, to better understand the results, we conducted a correlation analysis between the RT (mean of each individual’s median per conditions - as per the earlier analysis) and *b* values and between the RT and x_o_ values. While the RT/ x_o_ correlation yielded no significant results, the RT/*b* correlation did have some significant values (see figure 8). These correlations are exploratory and did not undergo any corrections (i.e. Bonferroni). To further interpret this result, we also conducted One-Sample Student T-tests between the PSS values and 0 (synchronous), which showed that only the Congruent/High condition (*p* = 0.001) was significantly different from 0, although the Incongruent/Low condition was closer to significance that the other non-significant categories (Congruent/Low, *p* = 0.790, Incongruent/High, *p* = 0.804, Incongruent/Low, *p* = 0.195) (see figure 6).

**Figure 8.**
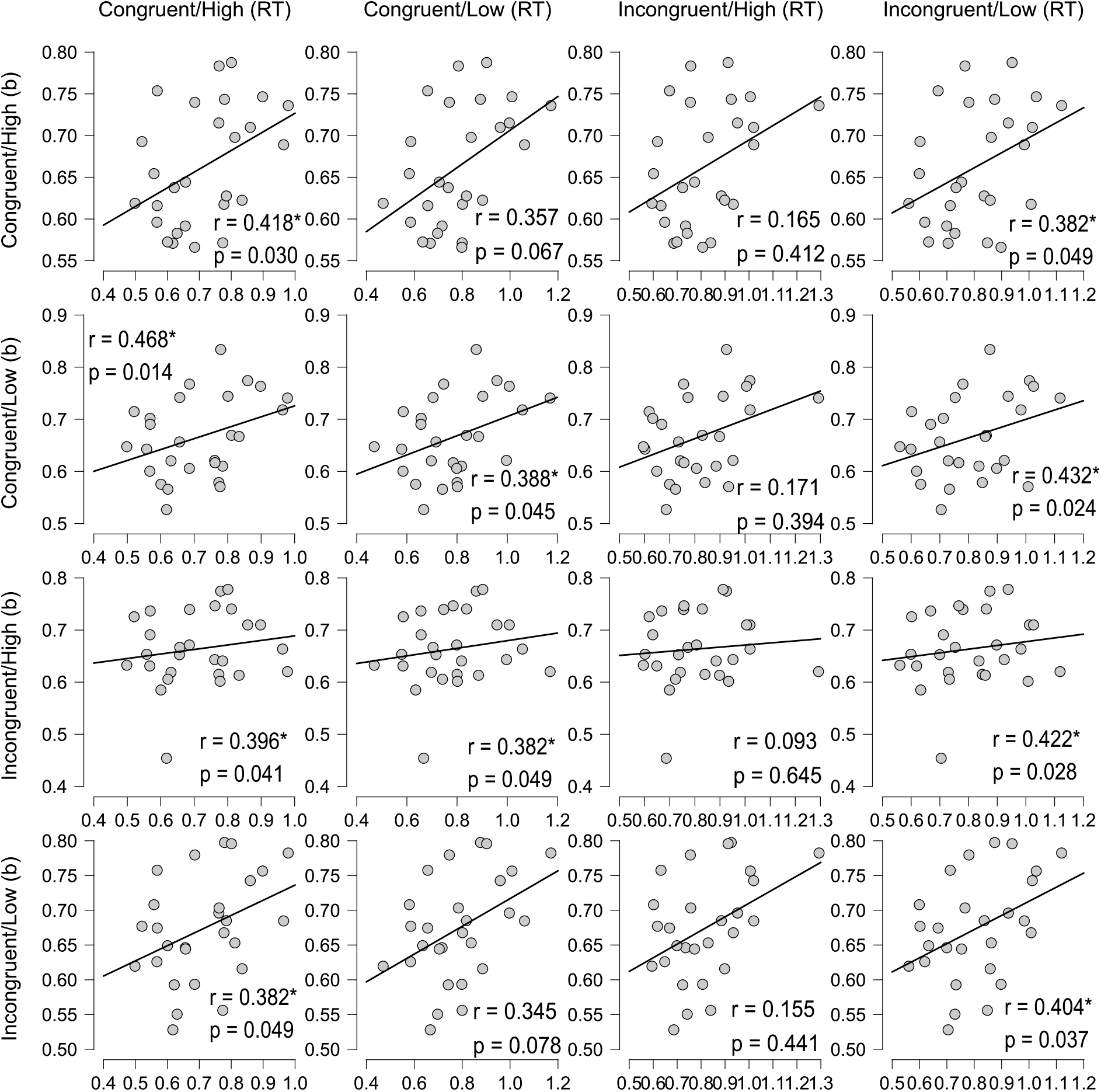
Results (plots, Pearson’s r values, and p-values) of the RT/*b* values correlation. An asterisk (*) indicates *p* < 0.05.

## Discussion

In this study, we used a novel paradigm that manipulated both semantic congruency and valence to measure the impact of these variables on the perceived timing of multisensory stimuli. We showed that: (1) as per hypothesis 1, congruent conditions had a main effect of significantly faster response times but unexpectedly, had no significant main effect on TOJ; (2) following the 2^nd^ hypothesis, valence did have a significant main effect on both RT and TOJ for the PSS values (high valence had a faster response time than low valence); and (3), with respect to the 3^rd^ hypothesis, valence and congruence did interact in both RT and TOJ. Specifically, for RT, high valence and congruent stimuli had the fastest response time (of all conditions) and the Congruent/Low condition had a significantly faster response time than Incongruent/High condition, and for TOJ, the Congruent/High stimuli had a significantly earlier PSS (requiring the sound to be presented before the image) than all other conditions.

The response time results were as expected and do support findings from the previous literature (Laurienti et al., 2004; Steinweg & Mast, 2017). Looking at semantic congruency, our results directly support the findings of Steinweg & Mast (2017) in that incongruent conditions have a slower response than congruent conditions. As per Steinweg & Mast (2017), we believe that this is most likely the result of an incongruity caution effect - participants require more time to make a response when presented with two incongruent stimuli. When the stimuli are congruent, there may be a multisensory integration and facilitation effect, while with incongruent stimuli this integration may not take place (Chen & Spence, 2010; Steinweg & Mast, 2017). This, combined with the incongruity caution effect, accounts for the main effect of congruence in our results. Our study also investigated the impact of valence on response time. There was both a main effect of valence on RT and a significant interaction effect between congruence and valence. This does follow our expected results and supports findings in previous affective priming studies that show that high valence may have a speeded response time (Yao et al., 2019; Yao & Wang, 2013). Previous use of valence in non-priming studies is sparse, thus the results from this study shed light on a very important yet understudied aspect of multisensory integration. As for the interaction effect, the congruent and high valence condition resulted in a faster RT than all of the other conditions, which also support previous literature on congruence and valence. Our results demonstrate that valence is an important part in studies and should be controlled for or evaluated in studies using images - it is important to consider, as it does impact results.

Moving to the temporal order judgement task, our results show that there is a main effect of valence, an interaction effect of valence and congruence on TOJ for the PSS values, but that there is no significant main effect of congruence. Looking first at the JND (*b*-value from the sigmoidal regression), our results (both the true values and the transformed values) show that there is no effect of congruence or valence. This supports Vatakis & Spence (2008) who also found no effect of congruency with non-speech video stimuli but opposes findings from Vatakis & Spence (2007) who found a lower JND (participants are better able to discriminate temporal order) for mismatched (incongruent) conditions of speech. Our results are perhaps different from Vatakis & Spence (2007) due to a difference in stimuli type. Both Vatakis & Spence (2007) and (2008) used video clips with accompanying sounds while we used static images, and Vatakis & Spence (2007) used speech stimuli, which is thought to have different integration qualities as compared to non-speech stimuli (Jones & Jarick, 2006). Thus, our results support some of the existing literature and provide further insight into the impact of congruence on static, non-speech stimuli. This area should be a focus of future research to further understand the impact of congruence on different multi-modal stimuli.

Looking now at the PSS (*x_0_* value from the regression), there were no main effects of congruency, however, there was a significant main effect of valence and a significant interaction effect (of nearly medium effect size (ɳ^2^= 0.05) and large effect size (ɳ^2^= 0.226) respectively) (Lakens, 2013). For the main effect of valence, in order for participants to perceive the stimuli as simultaneous, the sound was required to be presented significantly earlier for the high valance stimuli as compared to the low valence stimuli. There is very little literature available that investigates the impact of valence on TOJ tasks, thus this result provides a base for future research on this topic. Looking now at the interaction effect, compared to the Congruent/Low (0.005 s), Incongruent/High (−0.004 s), and Incongruent/Low (−0.027 s) condition, the Congruent/High condition (−0.051 s) required the sound to be presented significantly earlier than the image. This does, in part, support previous literature, albeit both Vatakis & Spence (2007) and (2008) reported on PSS primarily for the sake of completeness, not as indicative of a strong effect. Vatakis & Spence (2007) reported a necessary auditory lead for most of the congruent conditions (visual lead was required for the experiments with syllables and differing gender of speaker) and visual lead for the incongruent condition, while Vatakis & Spence (2008) require a visual lead for both congruent and incongruent conditions. A study looking at TOJ data for different stimuli found that a flash-click stimuli required the visual stimulus to precede audio, while a video stimuli needed the sound to precede video to have perceived simultaneity (Van Eijk et al., 2008). This shows that type of stimuli utilized in a study can impact the PSS thus potentially accounting for the variation between our results and other studies. The effect size of the interaction effect was large (ɳ^2^= 0.226), yet further research is required to further illuminate this issue.

Now looking at the exploratory correlation analyses, we find that there is a significant positive correlation between the TBW width (*b*-values) and RT for the categories that differed most from 0 in their PSS (Congruent/High significantly and Incongruent/Low nearing significance). As the correlations did not apply the Bonferroni correction, any conclusions should be considered with this in mind. While the “why” may not be evident from this, these results do illuminate this subject further. Future research should continue in this area.

Based on the current results, we find that, like Vatakis & Spence (2007), there is an effect of congruency on TOJ and, new to the field, that valence also has an impact. While the main effects may not yet be clear, the interaction effect and correlational effects lead us to believe that these factors do impact PSS, requiring further research.

Finally, this study presents a unique and relatively novel use of the online study environment for such behavioural studies. Crowdsourcing online platforms such as Mechanical Turk and Prolific Academic have been commonly used for studies involving surveys or other similar methods, but have only recently been used to investigate behavioural methods (such as RT and TOJ, as done here) using different software packages (Bazilinskyy & De Winter, 2018; Bridges et al., 2020; Woods et al., 2015). While limited in control of stimuli presentation, the online study environment opens up possibilities for a larger range and number of participants within a shorter period of time, ultimately greatly benefitting the research process. This study is one of the first to use this mode of investigation to measure RT and TOJ.

### Limitations

While we believe the methodology from this study is strong, there are some limitations to this design that should be considered. A first limitation is a possible limit on the scope of the design: this study only used two types of images and sounds (cats and snakes). It is possible that these are unique cases, and thus are not adequately representative of their categories as a whole. Additionally, we only included two levels of valence (high/low), thus, future research should expand the number of categories of valence (high/medium/low valence) as well as using more and different examples from each category for both images and sounds. Next, there are known to be differences in temporal order judgement (TOJ) tasks and simultaneity judgement (SJ) tasks, which this study did not evaluate due to feasibility constraints (Basharat et al., 2018; Van Eijk et al., 2008). Future research should investigate SJ/TOJ differences as it is a valuable vein of research that does help elucidate results. Finally, while the online environment provides unique advantages in terms of ease of recruitment it does have some limitations. This includes not being able to control the stimuli (screen size, brightness, distance, sound volume, clarity, etc.). We made all possible efforts to control these variables through instruction to participants but ultimately, we were unable to control this for each participant. Despite these limitations, this study provides a strong base and has the potential to guide future research in the understudied areas on congruence and valence impacting multisensory integration.

## Conclusion

This study has demonstrated that there are effects of both semantic congruency and valence on multisensory integration. Semantically congruent stimuli result in faster response times. High valence results in faster response times and an earlier PSS. High valence and congruent conditions result in the fastest response times and a significantly earlier PSS. RT and *b* values are correlated, and this correlation is moderated by the PSS (when the PSS was farther from 0 there was a stronger correlation between RT and *b*). This study provides new evidence that supports previous research on semantic congruency and presents a novel incorporation of valence into behavioural responses. Future research should attempt to expand the scope of design and incorporate an SJ task to further the understanding of this field.

## Acknowledgements

We would particularly like to thank Damien Masson for his assistance in coding the tasks and Rob McIlroy for his feedback throughout development of the study.

## References

Avero, P., & Calvo, M. G. (2006). Affective priming with pictures of emotional scenes: The role of perceptual similarity and category relatedness. Spanish Journal of Psychology, 9(1), 10–18. https://doi.org/10.1017/S1138741600005928

Barutchu, A., Spence, C., & Humphreys, G. W. (2018). Multisensory enhancement elicited by unconscious visual stimuli. Experimental Brain Research, 236(2), 409–417. https://doi.org/10.1007/s00221-017-5140-z

Basharat, A., Adams, M. S., Staines, W. R., & Barnett-Cowan, M. (2018). Simultaneity and Temporal Order Judgments Are Coded Differently and Change With Age: An Event-Related Potential Study. Frontiers in Integrative Neuroscience, 12, 15. https://doi.org/10.3389/fnint.2018.00015

Basharat, A., Mahoney, J. R., & Barnett-Cowan, M. (2019). Temporal Metrics of Multisensory Processing Change in the Elderly. Multisensory Research, 32(8), 715–744. https://doi.org/10.1163/22134808-20191458

Bazilinskyy, P., & De Winter, J. C. F. (2018). Crowdsourced Measurement of Reaction Times to Audiovisual Stimuli With Various Degrees of Asynchrony. Human Factors, 60(8), 1192–1206. https://doi.org/10.1177/0018720818787126

Bechlivanidis, C., & Lagnado, D. A. (2016). Time reordered: Causal perception guides the interpretation of temporal order. Cognition, 146, 58–66. https://doi.org/10.1016/j.cognition.2015.09.001

Bedard, G., & Barnett-Cowan, M. (2016). Impaired timing of audiovisual events in the elderly. Experimental Brain Research, 234(1), 331–340. https://doi.org/10.1007/s00221-015-4466-7

Box, G. E. P., & Cox, D. R. (1964). An Analysis of Transformations. Journal of the Royal Statistical Society: Series B (Methodological), 26(2), 211–243. https://doi.org/10.1111/j.2517-6161.1964.tb00553.x

Bradley, M. M., & Lang, P. J. (2017). International Affective Picture System. In V. Zeigler-Hill & T. K. Shackelford (Eds.), Encyclopedia of Personality and Individual Differences (pp. 1–4). Springer International Publishing. https://doi.org/10.1007/978-3-319-28099-8_42-1

Bridges, D., Pitiot, A., MacAskill, M. R., & Peirce, J. W. (2020). The timing mega-study: Comparing a range of experiment generators, both lab-based and online. PeerJ, 8. https://doi.org/10.7717/peerj.9414

Chen, Y. C., & Spence, C. (2010). When hearing the bark helps to identify the dog: Semantically-congruent sounds modulate the identification of masked pictures. Cognition, 114(3), 389–404. https://doi.org/10.1016/j.cognition.2009.10.012

Chen, Y. C., & Spence, C. (2017). Assessing the role of the “unity assumption” on multisensory integration: A review. In Frontiers in Psychology (Vol. 8, Issue MAR, p. 445). Frontiers Research Foundation. https://doi.org/10.3389/fpsyg.2017.00445

Colonius, H., & Diederich, A. (2017). Measuring multisensory integration: From reaction times to spike counts. Scientific Reports, 7(1). https://doi.org/10.1038/s41598-017-03219-5

Cox, D., & Hong, S. W. (2015). Semantic-based crossmodal processing during visual suppression. Frontiers in Psychology, 6(JUN), 722. https://doi.org/10.3389/fpsyg.2015.00722

Crouzet, S. M., Joubert, O. R., Thorpe, S. J., & Fabre-Thorpe, M. (2012). Animal Detection Precedes Access to Scene Category. PLoS ONE, 7(12), e51471. https://doi.org/10.1371/journal.pone.0051471

Crump, M. J. C., McDonnell, J. V., & Gureckis, T. M. (2013). Evaluating Amazon’s Mechanical Turk as a Tool for Experimental Behavioral Research. PLoS ONE, 8(3). https://doi.org/10.1371/journal.pone.0057410

Delong, P., & Noppeney, U. (2021). Semantic and spatial congruency mould audiovisual integration depending on perceptual awareness. Scientific Reports, 11(1), 10832. https://doi.org/10.1038/s41598-021-90183-w

Der, G., & Deary, I. J. (2017). The relationship between intelligence and reaction time varies with age: Results from three representative narrow-age age cohorts at 30, 50 and 69 years. Intelligence, 64, 89–97. https://doi.org/10.1016/j.intell.2017.08.001

Doehrmann, O., & Naumer, M. J. (2008). Semantics and the multisensory brain: How meaning modulates processes of audio-visual integration. In Brain Research (Vol. 1242, pp. 136–150). Elsevier. https://doi.org/10.1016/j.brainres.2008.03.071

Gerdes, A. B. M., Wieser, M. J., & Alpers, G. W. (2014). Emotional pictures and sounds: A review of multimodal interactions of emotion cues in multiple domains. In Frontiers in Psychology (Vol. 5, Issue DEC). Frontiers Research Foundation. https://doi.org/10.3389/fpsyg.2014.01351

Gottfried, J. A., & Dolan, R. J. (2003). The nose smells what the eye sees: Crossmodal visual facilitation of human olfactory perception. Neuron, 39(2), 375–386. https://doi.org/10.1016/S0896-6273(03)00392-1

Howard, I. P., & Templeton, W. B. (1966). Human spatial orientation. John Wiley & Sons.

Jones, J. A., & Jarick, M. (2006). Multisensory integration of speech signals: The relationship between space and time. Experimental Brain Research, 174(3), 588–594. https://doi.org/10.1007/s00221-006-0634-0

Kurdi, B., Lozano, S., & Banaji, M. R. (2017). Introducing the Open Affective Standardized Image Set (OASIS). Behavior Research Methods, 49(2), 457–470. https://doi.org/10.3758/s13428-016-0715-3

Lakens, D. (2013). Calculating and reporting effect sizes to facilitate cumulative science: a practical primer for t-tests and ANOVAs. Frontiers in Psychology, 4(NOV). https://doi.org/10.3389/FPSYG.2013.00863

Laurienti, P. J., Kraft, R. A., Maldjian, J. A., Burdette, J. H., & Wallace, M. T. (2004). Semantic congruence is a critical factor in multisensory behavioral performance. Experimental Brain Research, 158(4), 405–414. https://doi.org/10.1007/s00221-004-1913-2

Love, S. A., Petrini, K., Cheng, A., & Pollick, F. E. (2013). A Psychophysical Investigation of Differences between Synchrony and Temporal Order Judgments. PLoS ONE, 8(1). https://doi.org/10.1371/journal.pone.0054798

McGurk, H., & MacDonald, J. (1976). Hearing lips and seeing voices. Nature, 264(5588), 746–748. https://doi.org/10.1038/264746a0

Praß, M., Grimsen, C., König, M., & Fahle, M. (2013). Ultra Rapid Object Categorization: Effects of Level, Animacy and Context. PLoS ONE, 8(6). https://doi.org/10.1371/journal.pone.0068051

Schmidt, R. A. (1988). Motor control and learning: a behavioral emphasis (2nd ed.). Human Kinetics.

Spilcke-Liss, J., Zhu, J., Gluth, S., Spezio, M., & Gläscher, J. (2019). Semantic Incongruency Interferes With Endogenous Attention in Cross-Modal Integration of Semantically Congruent Objects. Frontiers in Integrative Neuroscience, 13, 53. https://doi.org/10.3389/fnint.2019.00053

Stein, B. E., Laurienti, P. J., Wallace, M. T., & Stanford, T. R. (2002). Multisensory Integration. In *Encyclopedia of the Human Brain* (pp. 227–241). Elsevier. https://doi.org/10.1016/b0-12-227210-2/00225-9

Stein, B. E., & Stanford, T. R. (2008). Multisensory integration: Current issues from the perspective of the single neuron. In Nature Reviews Neuroscience (Vol. 9, Issue 4, pp. 255–266). Nature Publishing Group. https://doi.org/10.1038/nrn2331

Steinweg, B., & Mast, F. W. (2017). Semantic incongruity influences response caution in audio-visual integration. Experimental Brain Research, 235(1), 349–363. https://doi.org/10.1007/s00221-016-4796-0

Thaler, L., Schütz, A. C., Goodale, M. A., & Gegenfurtner, K. R. (2013). What is the best fixation target? The effect of target shape on stability of fixational eye movements. Vision Research, 76, 31–42. https://doi.org/10.1016/j.visres.2012.10.012

Topolinski, S., & Deutsch, R. (2013). Phasic affective modulation of semantic priming. Journal of Experimental Psychology: Learning Memory and Cognition, 39(2), 414–436. https://doi.org/10.1037/a0028879

Van Eijk, R. L. J., Kohlrausch, A., Juola, J. F., & Van De Par, S. (2008). Audiovisual synchrony and temporal order judgments: Effects of experimental method and stimulus type. Perception and Psychophysics, 70(6), 955–968. https://doi.org/10.3758/PP.70.6.955

Vatakis, A., & Spence, C. (2007). Crossmodal binding: Evaluating the “unity assumption” using audiovisual speech stimuli. Perception and Psychophysics, 69(5), 744–756. https://doi.org/10.3758/BF03193776

Vatakis, A., & Spence, C. (2008). Evaluating the influence of the “unity assumption” on the temporal perception of realistic audiovisual stimuli. Acta Psychologica, 127(1), 12–23. https://doi.org/10.1016/j.actpsy.2006.12.002

Woods, A. T., Velasco, C., Levitan, C. A., Wan, X., & Spence, C. (2015). Conducting perception research over the internet: A tutorial review. In PeerJ (Vol. 2015, Issue 7). PeerJ Inc. https://doi.org/10.7717/peerj.1058

Yao, Z., & Wang, Z. (2013). The effects of the concreteness of differently valenced words on affective priming. Acta Psychologica, 143(3), 269–276. https://doi.org/10.1016/j.actpsy.2013.04.008

Yao, Z., Zhu, X., & Luo, W. (2019). Valence makes a stronger contribution than arousal to affective priming. PeerJ, 2019(10). https://doi.org/10.7717/peerj.7777

